# Why is *Babesia* not killed by artemisinin like *Plasmodium*?

**DOI:** 10.1101/2022.08.01.502256

**Authors:** Wenwen Si, Chuantao Fang, Chuang Liu, Meng Yin, Wenyue Xu, Yanna Li, Xiaoli Yan, Yujuan Shen, Jianping Cao, Jun Sun

**Author notes:** Corresponding author: Email address (JS). These authors contributed equally to this work (Wenwen Si, Chuantao Fang, Chuang Liu).

## Abstract

*Babesia* spp. are intraerythrocytic apicomplexan organisms digesting hemoglobin similar to intraerythrocytic *Plasmodium* spp. However, unlike *Plasmodium* spp., *Babesia* spp. are not sensitive to artemisinin, The difference between *Babesia* genomes and *Plasmodium* genomes revealed that smaller *Babesia* genomes lack numerous genes, especially haem synthesis-related genes. Single-cell sequencing analysis showed that different groups of *B. microti* with expressed pentose phosphate pathway (PPP)-related, DNA replication-related, antioxidation-related, glycolysis-related, and glutathione-related genes were not as sensitive to artemether as *P. yoelii 17XNL*. Especially, PPP-related, DNA replication-related, and glutathione-related genes were inactively expressed compared with *P. yoelii 17XNL*. Adding iron supply in vivo can promote the reproduction of *B. microti*. These results suggest that *Babesia* spp. lack a similar mechanism to that in malaria parasites, by which haem or iron in hemoglobin is utilized, but it likely leads to their insensitivity to artemisinin in turn.

**Author summary:** *Babesia* and *Plasmodium* are similar in many ways, from morphology to life history. In particular, both ingest and break down hemoglobin. However, compared with *Plasmodium*, *Babesia* cannot form hemozoin with less pathogenicity and insensitivity to artemisinin. Recent studies suggest that artemisinin can kill malaria parasites through free-radical and iron-capture effects, indicating that iron and haem play a key role in the sensitivity of malaria parasites to artemisinin. The *Babesia* genome is smaller and does not contain haem synthesis-related genes, indicating low requirements and utilization of haem and iron (HI). Moreover, we found that the expression of PPP-related and DNA replication-related genes is not active, distinctly different from malaria parasites. However, adding iron supply in vivo can increase the infection rate of *B. microti*. Therefore, we hypothesized that *Babesia* lacks mechanisms for the efficient utilization of HI, resulting in low requirements for HI, and therefore insensitivity to artemisinin.

## Introduction

*Babesia* is a protozoan parasite that leads to a hemolytic disease known as babesiosis. More than 100 species of *Babesia* have been identified, and a few species have been documented as pathogenic in humans [1]. Similarly, hundreds of types of *Plasmodium* species have been identified, and five of them cause malarial disease in humans. Both are intraerythrocytic protozoa, resulting in common clinical signs such as hemolytic anemia, fever, choloplania, hemoglobinuria, and an enlarged spleen[2]. The life cycle of both species involves two hosts: one is a vertebrate; another is an insect. During a blood meal, both can introduce sporozoites into the vertebrate host, where they can infect erythrocytes and undergo asexual reproduction. Afterward, both differentiate into male and female gametocytes. Then, they are ingested by the insect host, where gametes unite and undergo a sporogonic cycle, resulting in sporozoites. When the insect hosts bite the next vertebrate hosts, another new cycle begins. These two intraerythrocytic protozoan parasites share so similar life cycle, morphology, pathogenicity, and vertebrate and insect vectors that they are often confused.

Interestingly, artemisinin and its derivatives (ARTs) efficiently kill malaria parasites but only slightly inhibit the growth of *Babesia* species [3–5]. Although they both digest hemoglobin for the supplement of amino acids, only *Plasmodium* produces hemozoin in the metabolism processes. Previously, hemozoin was widely accepted as an insoluble metabolic byproduct of hemoglobin digestion, without biological function during parasites infecting red blood cells (RBC). However, increasing evidences support that hemozoin is considered a critical vector of haem and iron (HI) and is responsible for storing and utilizing iron [6–8]. In particular, malaria parasites always maintain a high haem level (1.6 µmol/L) throughout their development in RBCs, far more than other parasites [9]. Meanwhile, amounts of hemozoin are transferred to gametocytes, suggesting a biological significance of hemozoin in the reproduction of *Plasmodium*. More interestingly, malaria parasites lacking the formation of hemozoin show similar phenotypes to *Babesia* in morphology and pathogenicity with less virulence and reproduction, such as chloroquine-resistant parasites [10, 11]. It is not clear why *Babesia* does not produce hemozoin, but it is very clear that hemozoin is a haem polymer, naturally storing a large amount of HI. Malaria parasites are sensitive to iron chelators, suggesting that they depend on iron more than other organisms that are not sensitive to iron chelators. In particular, recent research proposed that artemisinin kills *Plasmodium* through the iron-capture effect[12]. Apparently, *Plasmodium* and *Babesia* have different HI requirements, which may be relevant to their different sensitivity to ARTs. Why *Babesia* do not require or store the high amount of HI as *Plasmodium* did, why *Babesia* do not produce hemozoin like *Plasmodium*, and whether different HI requirements determine their different fecundity remain to be elucidated. To investigate the underlying mechanism, we compared the genomes of these two species and analyzed the effect of artemether action on *P. yoelii* 17XNL and *B. microti* using single–cell transcriptomic sequencing, and the effect of adding iron supply on *Babesia* growth in vivo.

## Materials and methods

### Ethics statement

This study was carried out in strict accordance with the recommendations of the Regulations for the Administration of Affairs Concerning Experimental Animals of the State Science and Technology Commission. The protocol was approved by the Internal Review Board of Tongji University School of Medicine (TJLAC-017–039).

### Comparison between *Babesia* and *Plasmodium* genomes

To systematically explore the distinction between the genomes of 53 *Plasmodium* and 6 *Babesia* (S1 Table), *Plasmodium* Informatics Resource database (PlasmoDB), Prioplasma Informatics Resources database (Piroplasmadb), and Eukaryotic Pathogen, Vector and Host Informatics Resource (VEuPathDB) [13], were utilized to classify genes that have a specified orthology-based phylogenetic profile into three subgroups. To further explore the functions of these three categories of genes, Gene Ontology (GO) enrichment and metabolic pathway enrichment (using the algorithms from the Kyoto Encyclopedia of Genes and Genomes and Metabolic Pathways From all Domains of Life pathway database) were carried out using the online tools on PlasmoDB (https://plasmodb.org/plasmo/app/workspace/strategies/), with a default p-value cutoff of 0.05.

### Experimental animals

*Plasmodium yoelii* 17XNL-EGFP was provided by Dr. Ana Rodriguez (New York University) and Dr. Wenyue Xu (Department of Pathogenic Biology, Army Medical University). The *Babesia microti* strain ATCC®PRA-99TM was provided by the National Institute of Parasitic Diseases, Chinese Center for Disease Control and Prevention. *Plasmodium yoelii* 17XNL-EGFP were cultured in female Balb/c mice or ICR mice. *Babesia microti* were cultured in NOD/SCID mice. All mice were purchased from Shanghai SLAC Laboratory Animal Co. Ltd. (Shanghai, China) and stored in the Laboratory Animal Center of Tongji University. The NOD/SCID and Balb/c mice were inoculated with an intraperitoneal injection of 200 ul of the cell suspension, which contained 1-5×10^7^ *B.microti* or *P. yoelii* 17XNL-EGFP. When the infection rate of *P. yoelii 17XNL* reached 25%-50 %, artemether (100 mg/kg) was fed to the mice. Then, blood samples were collected at 0 and 24 h after artemether treatment, respectively.

### Cells isolation and cell sorting

At 0 and 24 h after artemether treatment, 1-2 drops of the blood sample from the mouse infected with *P. yoelii* 17XNL-EGFP parasites were collected in 10 ml RPMI 1640 media in a 15 ml tube and immediately transferred onto ice, along with the controls for cell sorting. Cell samples were sorted on a BD FACS-Aria II by using UV, blue, and red lasers at 355, 488, and 633 nm, respectively. After sorting, only infected erythrocytes were collected for the scRNA experiment, in accordance with a previous study [12]. Differently, 1-2 drops of the blood sample from the mouse infected with *B.microti* parasites were collected in 10 ml RPMI 1640 media with 5% serum. Then, they were stained with CD45 antibody in the dark at room temperature for 20 min, and incubated with Hoechst 33342 at room temperature for 10 min. The control group did not undergo any treatment. The samples were washed and resuspended in RPMI 1640 medium before proceeding to FACS sorting. According to the requirement of single-cell sequencing, BD FACS Aria II was used to obtain enough iRBCs. Positive cells were collected in RPMI 1640 medium supplemented with 5% serum. Cells were stained with 0.4% Trypan blue to check the viability on Countess® II Automated Cell Counter. Cell samples were sorted on a BD FACSAria II by using UV, blue and red lasers at 355, 488, and 633 nm, respectively. After sorting, only infected erythrocytes were collected for the scRNA experiment.

### Chromium 10X Genomics library and sequencing

Single-cell suspensions were loaded to 10X Chromium to capture approximately 3000 - 10,000 single cells according to the manufacturer’s instructions of the 10X Genomics Chromium Single-Cell 3’ kit. The following cDNA amplification and library construction steps were performed according to the standard protocol. Libraries were sequenced on an Illumina NovaSeq 6000 sequencing system (paired-end multiplexing run, 150bp) by Majorbio Co., Ltd. (Shanghai, China).

### scRNA-seq

The reads were processed using the Cell Ranger 4.0 pipeline with default and recommended parameters. FASTQs generated from Illumina sequencing output were aligned to the mouse genome, version GRCm38, using the STAR algorithm[14]. Next, gene-barcode matrices were generated for each individual sample by counting unique molecular identifiers and filtering non-cell associated barcodes. Finally, a gene-barcode matrix containing the barcoded cells and gene expression counts was generated. This output was then imported into the Seurat (v3.2.0) R toolkit for quality control and downstream analysis of the single cell RNA-seq data[15]. All functions were run with default parameters unless specified otherwise. The matrices to exclude low-quality cells were filtered using a standard panel of three quality criteria: (1) number of detected transcripts (number of unique molecular identifiers), (2) detected genes, and (3) percent of reads mapping to mitochondrial genes (quartile threshold screening criteria). The normalized data (NormalizeData function in Seuratpackage) were used for extracting a subset of variable genes. Variable genes were identified while controlling for the strong relationship between variability and average expression.

### Identification of cell types and subtypes by dimensional reduction and cluster analysis

Gene expressions from each voxel were normalized by the sctransform[16] in Seurat, which uses regularized negative binomial models to account for technical artifacts while preserving biological variance. Then, the top 30 principal components were calculated and used to construct the KNN graph. The Louvain algo-rithm was used to cluster the voxels. We visualized the clusters on a 2D map produced with Uniform Manifold Approximation and Projection (UMAP). For each cluster, we used the Wilcoxon rank-sum test to find significant deferentially expressed genes comparing the remaining clusters. SingleR was used to identify the cell type[17].

### Differential expression analysis and functional enrichment

The differential expression genes (DEGs) between two different samples or clusters were obtained using the function FindMarkers in Seurat, using a likelihood ratio test. Essentially, DEGs with |log2FC|>0.25 and Q value <= 0.05 were considered to be significantly different expressed genes. In addition, GO functional enrichment analysis was performed to identify which DEGs were significantly enriched in GO terms and metabolic pathways at Bonferroni-corrected p-value ≤0.05 compared with the whole-transcriptome background. GO functional enrichment analyses were carried out by Goatools (https://github.com/tanghaibao/Goatools).

### RNA velocity analysis

The RNA velocity was calculated based on the spliced and unspliced counts as previously reported [18], and cells that were present in the pseudotemporal ordering were used for the analysis. We estimated the RNA velocity by scVelo (https://scvelo.org) [19], a method of developmental trajectory analysis. It estimated the variation of RNA abundance over time by calculating the ratio of mRNA before and after splicing in cells and inferred the next possible differentiation direction of cells. To plot individual cell velocities, the UMAP and T-SNE embeddings in Seurat were exported.

### Iron dextran assay in vivo

In the in vivo experiment, 45 Balb/c mice were infected with (1-10) × 10^6^ *B. microti*-parasitized erythrocytes by intraperitoneal injection. Then, they were randomly divided into three groups. From the second day of infection, groups 1 and 2 were administered subcutaneous injections of 1 and 1.25 g/kg iron dextran every two days, respectively. By contrast, group 3 was injected subcutaneously with an equal amount of saline. Blood samples were then collected from their tail veins per day to examine the infection rates by Giemsa staining. Statistical analysis was conducted using a T-test (unpaired) with GraphPad Prism 8.0 software. A p-value less than 0.01 was considered statistically significant.

### Tissue preparation and immunofluorescence assay

Spleens were removed from the mice 12 days post-infection of *B.microti*. Then, they were fixed in a 10% neutral paraformaldehyde fix solution. Immunofluorescence samples were prepared and operated according to the protocol (see S1 File).

### Serum cytokine analysis

Multiplex kits for measuring cytokines were purchased from Bio-Rad (Bio-Plex Pro Mouse Cytokine Grp I Panel 23-plex). Cytokine analyses were performed by Wayen Biotechnologies (Shanghai, China) using the Bio-Plex MagPix System (Luminex, Austin, TX, USA) following the manufacturer’s instructions for the Luminex xMAP technology with multiplex beads. Bio-Plex Manager version 6.1 software (Luminex, Austin, TX, USA) was used to calculate cytokine concentrations among the uninfected normal group, infected group without iron dextran treatment, and infected group with 1.25 g/kg of iron dextran treatment group. A nonlinear least-squares minimization algorithm generated a curve fitted by a five-parameter logistic equation and determined the high and low limits of detection. Twenty-three cytokines were measured. The results are expressed as picograms per milliliter.

## Results

### Smaller *Babesia* genomes lack complete haem synthesis enzymes system compared with *Plasmodium* genomes

*Babesia* genomes contain four chromosomes. Their size range from 6 Mb to 15 Mb, such as *B. microti* (6.44 Mb), *B. bovis* (8.18 Mb), *B.bigemina* (12.84 Mb), *B. ovata* (14.45 Mb), *B.ovis* (8.38 Mb) and *B.divergens* (9.65 Mb) (S1 Table and Fig 1A). By contrast, the *Plasmodium* genomes contain about 14 chromosomes. Their size range from 14 to 38Mb, such as *P. falciparum* (23.49Mb), *P. vivax* (29.04Mb), *P. yoelii* (22.45 Mb), and *P. malariae* (31.92 Mb) (S1 Table and Fig 1A). To investigate the commonality and individuality between *Babesia* and *Plasmodium*, we systematically explored the distinction between the genomes of 53 *Plasmodium* and 6 *Babesia* (Fig. 1B). PlasmoDB, a well-known *Plasmodium* informatics resource[13], was utilized to classify genes that have a specified orthology-based phylogenetic profile into three subgroups: 669 *Plasmodium*-specific (S2 Table), 924 *Babesia*-specific (S3 Table), and 1591 *Plasmodium*-*Babesia*-shared genes (S4 Table). After GO (gene ontology) analysis, our results highlighted three terms: microtubule-based movement, fatty acid biosynthetic/metabolic process, and haem biosynthetic/metabolic process (Fig 1B). To reveal the difference of their core function, we compared their intersections and obtained another three subgroups: 1185 *Plasmodium*-specific (S5 Table), 453 *Babesia*-specific (S6 Table), and 1075 *Plasmodium-Babesia*-shared genes (S7 Table). Of note, there are some biological processes only in *Plasmodium*-specific genes group, such as porphyrin-containing compound biosynthetic process, haem biosynthetic process, porphyrin-containing compound metabolic process, tetrapyrrole biosynthetic process, tetrapyrrole metabolic process, protoporphyrinogen IX metabolic process, protoporphyrinogen IX biosynthetic process, haem metabolic process, pigment biosynthetic process, and pigment metabolic process (S2 and S5 Table). In particular, haem-synthesis genes widely exist in various organisms and play a vital role in the life process. We analyzed haem synthesis genes in *Plasmodium* and *Babesia* genomes.

**Fig 1.**
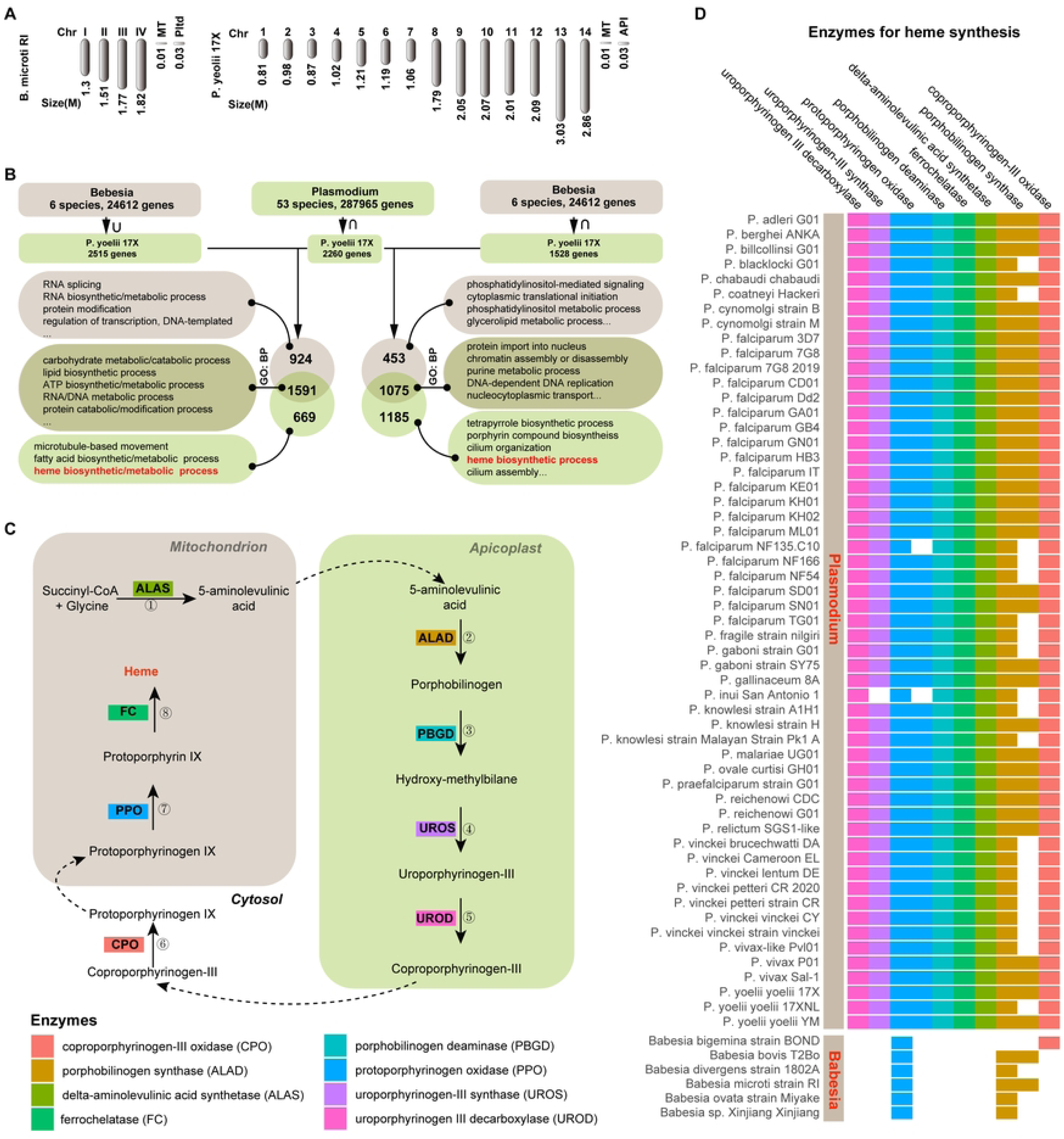
Comparison of *Babesia* and *Plasmodium* genomes and the distribution of haem synthesis-related genes in these genomes. **(A)** Schematic of *B. microti* and *P. yoelii* chromosomes. **(B)** Schematic of the analysis of the common and special genes of *B. microti* and *P. yoelii*. **(C)** Process of haem synthesis and eight key enzymes. (**D**) Distribution of haem synthesis-related genes in *Babesia* and *Plasmodium* genomes.

Haem is biologically synthesized by a complex and successive enzymatic reaction mediated by eight enzymes including coproporphyrinogen-III oxidase (CPO), delta-aminolevulinic acid dehydratase (ALAD), delta-aminolevulinic acid synthase (ALAS), ferrochelatase (FC), porphobilinogen deaminase (PBGD), protoporphyrinogen oxidase (PPO), uroporphyrinogen III decarboxylase (UROS), and uroporphyrinogen III synthase (UROD) (Fig 1C) [20]. To further evaluate the significance of haem metabolism, we analyzed the integrity of the haem biosynthesis pathway in 53 Plasmodium and 6 *Babesia* species in detail by using VEuPathDB. Approximately 92.3% (51/53) of *Plasmodium* species, have an integrated enzymatic system of de novo synthesis of haem, except *P. inui* San Antonio 1 and *P. falciparum* NF135.C10) (Fig 1D). However, only PPO and ALAD were identified in the genomes of six *Babesia* species (Fig 1D), suggesting that *Babesia* species lost the ability to biologically synthesize haem de novo. The results revealed that *Babesia* spp. cannot complete haem synthesis alone.

### Key enzyme genes in the pentose phosphate pathway (PPP) in *Babesia* are not as actively expressed as those in *Plasmodium*

In malaria parasites, hemoglobin is degraded to release haem. HI play a crucial role in connecting hemoglobin degradation with PPP to continuously produce nicotinamide adenine dinucleotide phosphate (NADPH) for the reduction of oxidized glutathione and thioredoxin systems and ribose-5-phosphate for nucleotide biosynthesis. This cycle is referred to as the “ Hemoglobin-Haem-Iron-PPP (HHIP) ” [12]. When the HHIP is sustained, DNA synthesis can continuously acquire ribose-5-phosphate[12]. We found that both parasites have complete PPP-related enzymes(Fig 2). Based on the distribution of parasites that expressed PPP-related genes in the UMAP plots, we found that these *Babesia* genes were not as actively expressed as those of *Plasmodium*. According to RNA velocity analysis (Figs 2A and 2B), enzyme genes, such as glucose-6-phosphate dehydrogenase-6-phosphogluconolactonase (PY17X_1321300), 6-phosphogluconate dehydrogenase, decarboxylating (PY17X_1322200) and transketolase (PY17X_0110700) were highly expressed at the early and middle stages in malaria parasites (Fig 2E). Similarly, these enzymes in *Babesia* are also expressed at these stages, but they are not as actively expressed as those in *Plasmodium* (Figs 2C - 2E).

**Fig 2.**
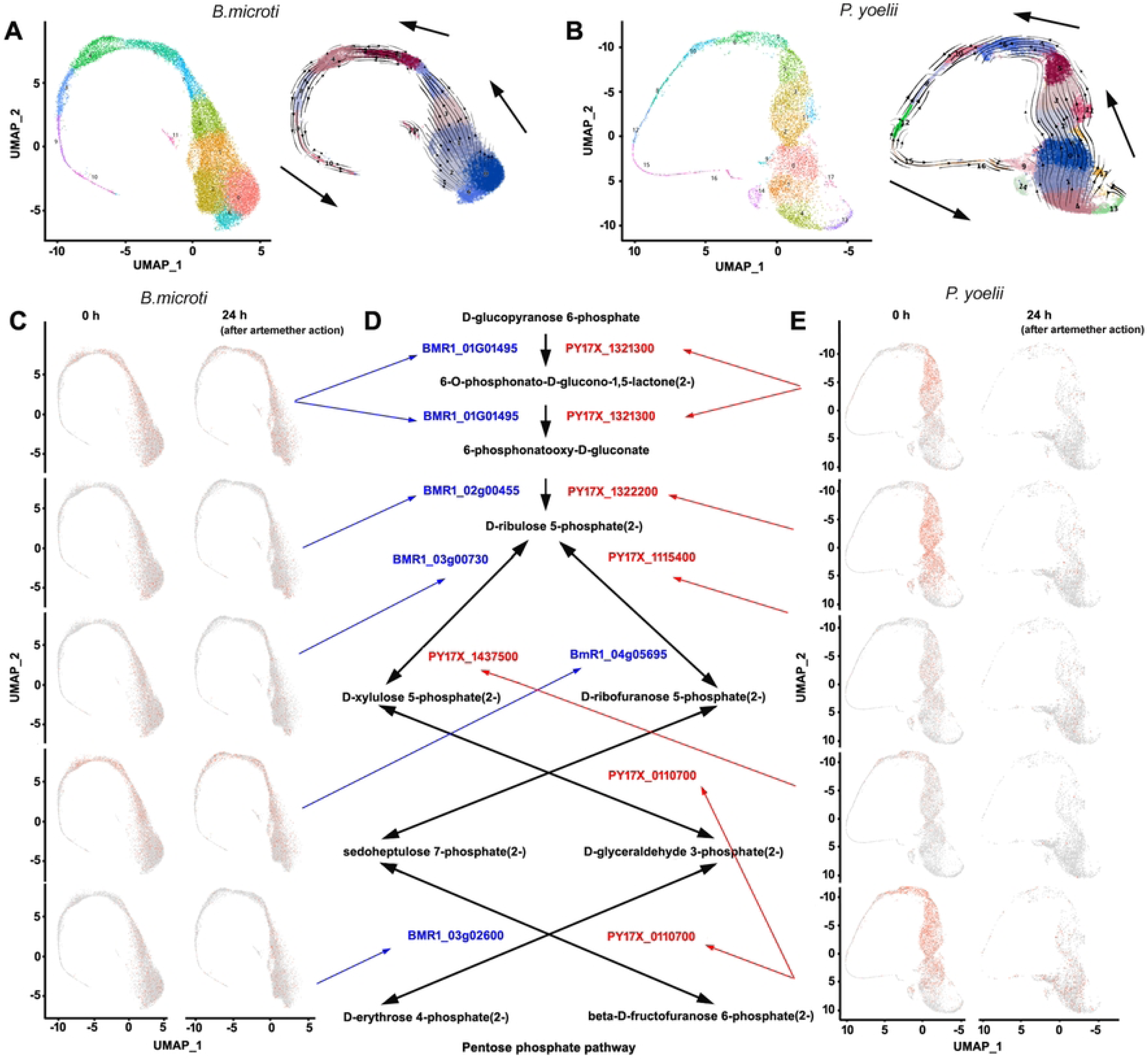
UMAP plots showing the expression characteristics of key enzyme genes in PPP in *B. microti* and *P. yoelii* 17XNL, and the sensitivity of these parasites that expressed these genes to artemether. **(A)** UMAP plot of single-cell transcriptomes (SCTs) from the artemether-treated and control groups of *B. microti.* RNA velocity analysis showing the developmental relationships among different clusters of *B. microti.* **(B)** UMAP plot of SCTs from the artemether-treated and control groups of *P. yoelii* 17XNL. RNA velocity analysis showing the developmental relationships among different clusters of *P. yoelii* 17XNL. **(C)** UMAP plots showing the expression characteristics of 6-phosphogluconolactonase (BMR1_01G01495), and 6-phosphogluconate dehydrogenase (BMR1_02g00455), ribulose-phosphate 3-epimerase (BMR1_03g00730), ribose 5-phosphate isomerase A (BmR1_04g05695), and transaldolase (BMR1_03g02600) and the sensitivity of these parasites that expressed these genes to 24 h of artemether treatment. **(D)** Schematic of PPP. **(E)** UMAP plots showing the expression characteristics of glucose-6-phosphate dehydrogenase-6-phosphogluconolactonase (PY17X_1321300), 6-phosphogluconate dehydrogenase, decarboxylating (PY17X_1322200), ribulose-phosphate 3-epimerase (PY17X_1437500), ribose-5-phosphate isomerase (PY17X_1115400), and transketolase (PY17X_0110700) and the sensitivity of these parasites that expressed these genes to 24 h of artemether treatment.

### *B. microti* that expressed PPP - related genes is not susceptible to artemether

Based on RNA velocity analysis, in the UMAP plot of *B. microti*, clusters 0 and 6 are the earliest development stage, whereas cluster 10 is the final stage (Fig 2A). In the UMAP plot of *P. yoelii,* clusters 1 and 4 are the earliest development stage, whereas cluster 16 is the final stage (Fig 2B).

The parasites that expressed key genes in PPP such as 6-phosphogluconolactonase (BMR1_01G01495), 6-phosphogluconate dehydrogenase (BMR1_02g00455), ribulose-phosphate 3-epimerase (BMR1_03g00730), ribose 5-phosphate isomerase A (BmR1_04g05695), and transaldolase (BMR1_03g02600) were hardly affected 24 h post-artemether treatment (Figs 2C and 2D). By contrast, most malaria parasites that expressed glucose-6-phosphate dehydrogenase-6-phosphogluconolactonase (PY17X_1321300), 6-phosphogluconate dehydrogenase, decarboxylating (PY17X_1322200), ribulose-phosphate 3-epimerase (PY17X_1437500), ribose-5-phosphate isomerase (PY17X_1115400), and transketolase (PY17X_0110700) were eliminated 24 h post-artemether treatment (Fig 2E).

### *B. microti* that expressed DNA synthesis-, antioxidation-, and glycolysis-related genes are not susceptible to artemether

In *B. microti* groups, most parasites that highly expressed DNA-related genes, such as DNA polymerase epsilon catalytic subunit 1 and DNA polymerase alpha catalytic subunit A, were not affected by artemether treatment. However, in *P. yoelii* 17XNL groups, most parasites that expressed similar genes, were eliminated 24 h post-artemether treatment(Figs 3A - 3D and S1 Fig.). The same thing also occurred in other *B. microti* parasite groups, for example, *B. microti*, that expressed antioxidation-related genes, such as peroxiredoxin and thioredoxin reductase, and glycolysis-related genes, such as pyruvate kinase and hexokinase (Figs 3E and 3G). In *P. yoelii* 17XNL groups, most of those that expressed similar genes were eliminated 24 h post-artemether treatment (Figs 3F and 3H). In addition, the *B. microti* at the late stage, which expressed merozoite trap-like protein and apical merozoite protein genes, were not sensitive to artemether either, unlike *P. yoelii* 17XNL (Figs 3I and 3J). Based on the results, nearly all stages in *B.microti*, involving DNA synthesis, antioxidation, glycolysis, and reproduction, are not susceptible to artemether, distinctly different *P. yoelii* 17XNL.

**Fig 3.**
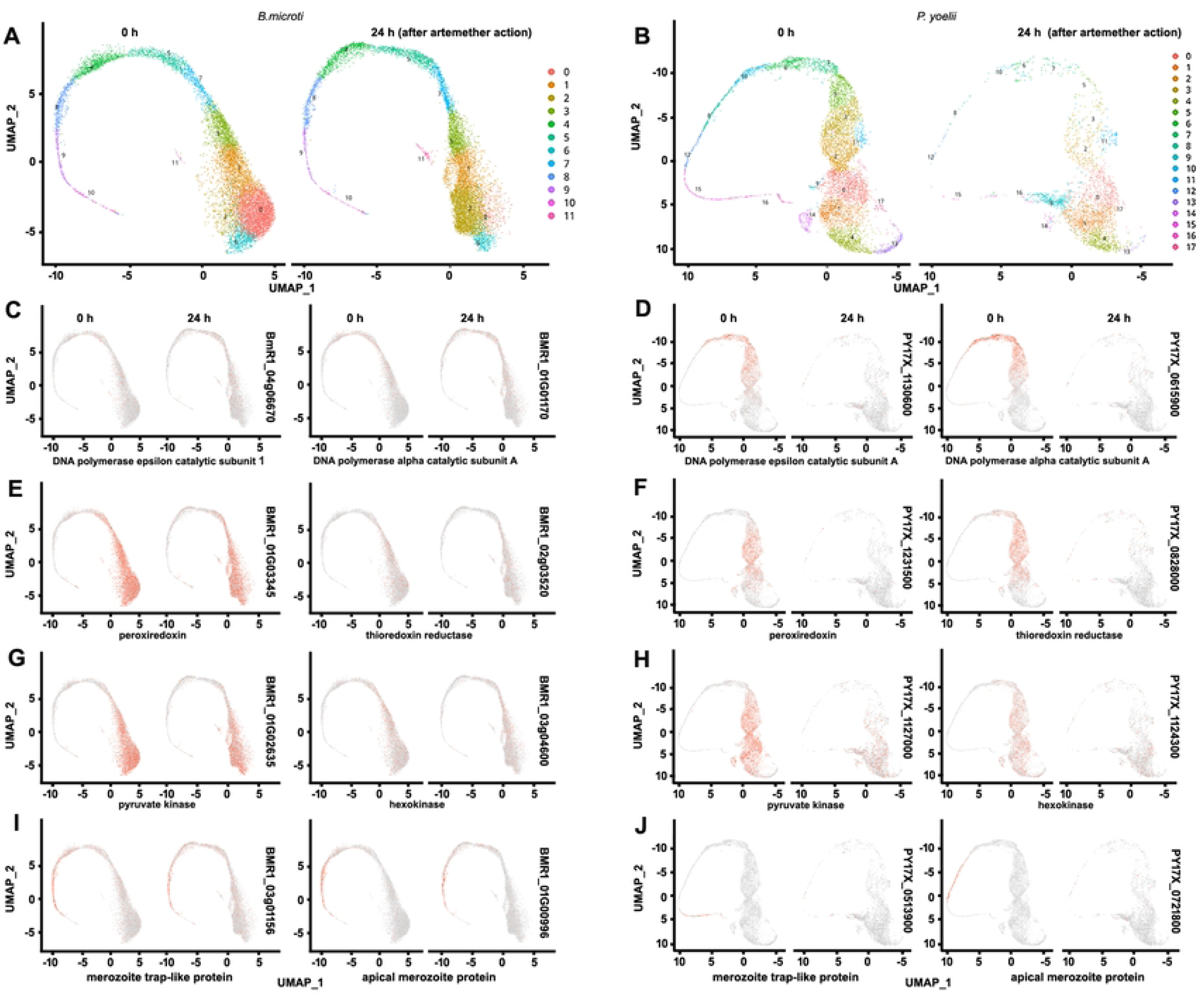
UMAP plots showing the sensitivity of *B. microti* and *P. yoelii* 17XNL that expressed different genes to 24 h of artemether treatment. **(A)** UMAP plot of single-cell transcriptomes (SCTs) from the artemether-treated and control groups of *B. microti*, showing the sensitivity of *B. microti* to 24 h of artemether treatment. **(B)** UMAP plot of SCTs from the artemether-treated and control groups of *P. yoelii* 17XNL, showing the sensitivity of *P. yoelii* 17XNL to 24 h of artemether treatment. **(C)** UMAP plots showing the sensitivity of *B. microti* that expressed DNA polymerase epsilon subunit 1 (BMR1_04g06670) and DNA polymerase alpha subunit A (BMR1_01g01170) to 24 h of artemether treatment. **(D)** UMAP plots showing the sensitivity of *P. yoelii* 17XNL that expressed DNA polymerase epsilon subunit A (PY17X_1130600) and DNA polymerase alpha subunit A (PY17X_0615900) to 24 h of artemether treatment. **(E)** Peroxiredoxin (BMR1_01g03345) and thioredoxin reductase (BMR1_02g03520). **(F)** Peroxiredoxin (PY17X_1231500) and thioredoxin reductase (PY17X_0828000). **(G)** Pyruvate kinase (BMR1_01g02635) and hexokinase (BMR1_03g04600).**(H)** Pyruvate kinase (PY17X_1127000) and hexokinase (PY17X_1124300). **(I)** Merozoite trap-like protein (BMR1_03g01156) and apical merozoite protein (BMR1_03g00996). **(J)** Merozoite trap-like protein (PY17X_0513900) and apical merozoite protein (PY17X_0721800).

### Glutathione-related genes and superoxide dismutase [Fe] gene of *B. microti* are not as actively expressed as those of *P. yoelii* 17XNL

In malaria parasites, hemoglobin is ingested and degraded, releasing haem. Haem is degraded by glutathione into iron and superoxide anion[21]. Superoxide dismutase [Fe] (SOD) can catalyze the dismutation of superoxide anion to hydrogen peroxide. Of note, in the HHIP cycle of malaria parasites, the degradation of haem by glutathione to iron is a key step [12]. The released iron will activate PPP to continuously produce NADPH and ribose-5-phosphate for nucleotide biosynthesis. We found that the number of *B. microti* that expressed glutathione synthase and thioredoxin / glutathione reductase genes was less than that of *P. yoelii 17XNL.* Moreover, the *B. microti* genome does not contain glutathione S-transferase gene, unlike *P. yoelii 17XNL* (Fig 4A). In addition, the expression of SOD gene of *B. microti* nearly involved all stages, rather than being concentrated in some specific stages, like *P. yoelii 17XNL* (Fig 4B). Furthermore, the number of *B. microti* that expressed the SOD gene at earlier stages was far less than that of *P. yoelii 17XNL*. Most of *P. yoelii 17XNL* that expressed these genes were eliminated 24 h of artemether treatment. By contrast, these parasites in *B. microti* were hardly affected by the same treatment (Figs 4A and 4B).

**Fig 4.**
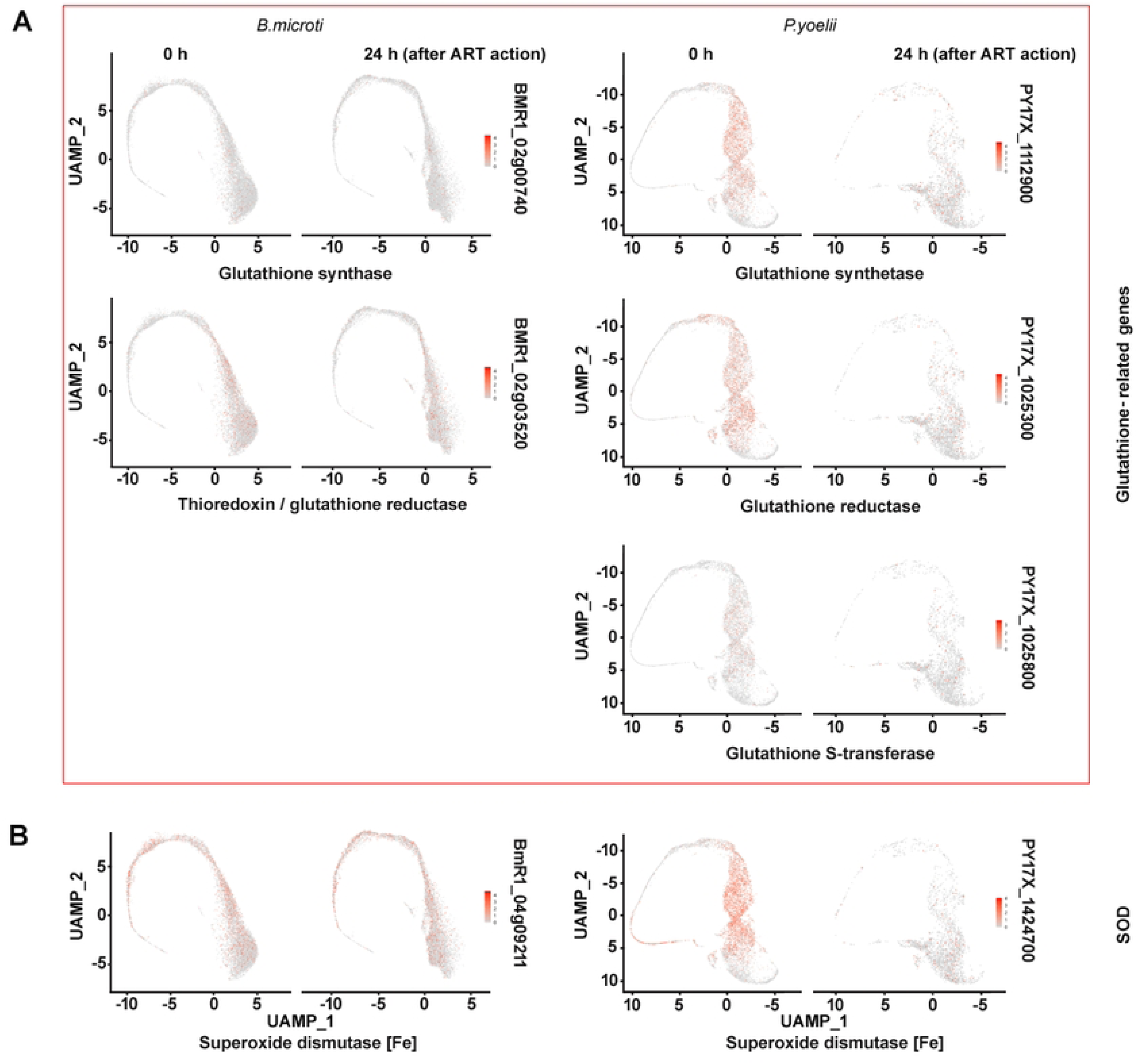
UMAP plots showing the differences in the number and gene expression activity of *B. microti* and *P. yoelii* 17XNL that expressed glutathione-related genes and superoxide dismutase gene. **(A)** UMAP plots showing the differences in the number and gene expression activity and artemether sensitivity of *B. microti* and *P. yoelii* 17XNL that expressed glutathione-related genes. Most of *P. yoelii* 17XNL that expressed these genes were eliminated 24 h of artemether treatment, whereas *B. microti* were hardly affected by the same treatment. **(B)** Differences in the number and gene expression activity and artemether sensitivity of *B. microti* and *P. yoelii* 17XNL that expressed superoxide dismutase [Fe] (SOD) gene.

### Adding iron supply promotes the reproduction of *P. yoelii* 17XNL

After mice were infected by *B. microti*, the infection rate peaked generally 8-10 days post-infection. When the mice were injected with iron dextran, the infection rate will increase gradually with the increase of time and dosage (Figs 5A, 5D and 5E). The result showed that the infection rate peaked 9 days post-infection. The infection rate in the experimental group was higher than that in the control group. Especially, the infection rate was significantly higher in the group that was administered subcutaneous injections of 1.25 g/kg iron dextran than in other groups (Figs 5A, 5D, 5E and 5F). All groups were infected by *B. microti*, and enlarged spleens could be observed (Fig 5B). There was no significant difference in the spleen weight except that the color of the spleen was darker in the experimental groups, especially in the group treated with 1.25g /kg iron dextran (Fig 5C).

**Fig 5.**
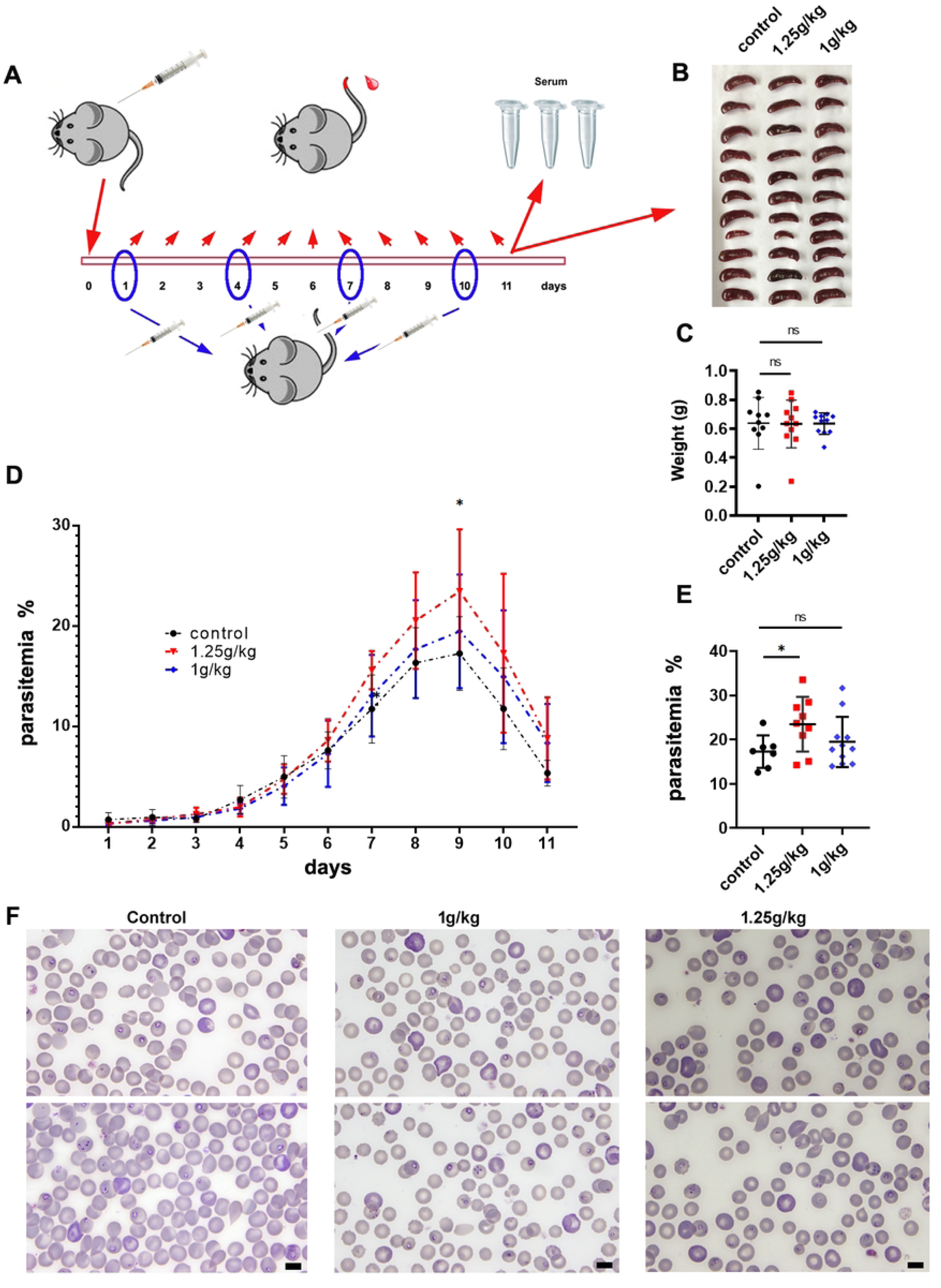
Effect of adding iron supply on the infection rate of *B.microti* in vivo. **(A)** Schematic of the experimental process. **(B)** Spleen samples from the infected mice in the control, 1 g/kg, and 1.25 g/kg test groups. **(C)** Comparison of the weight of spleen samples. **(D)** Effect of adding iron supply on the parasitemia of *B. microti*. **(E)** Comparison of the infection rates of *B.microti* on day 9. “ns” indicates not significant (p > 0.05), whereas the asterisk indicates a significant difference (*, p < 0.05). **(F)** Blood smears showing the morphology of *B. microti* and different infection rates in three groups on day 9. Scale bars indicate 5 μm.

### Macrophage M1/M2 polarization after iron supply in infected mice by *B. microti*

We identified the phenotype and number of macrophages using immunohistochemistry and fluorescence in situ hybridization in mouse spleen specimens. By detecting the changes in the fluorescence intensity of fluorescent antibodies of inducible nitric oxide synthase (iNOS) and CD206, we found that the number of macrophages positive for iNOS and CD206 is similar in all test groups and control groups. No significant changes were found among groups (Fig 6A).

**Fig 6.**
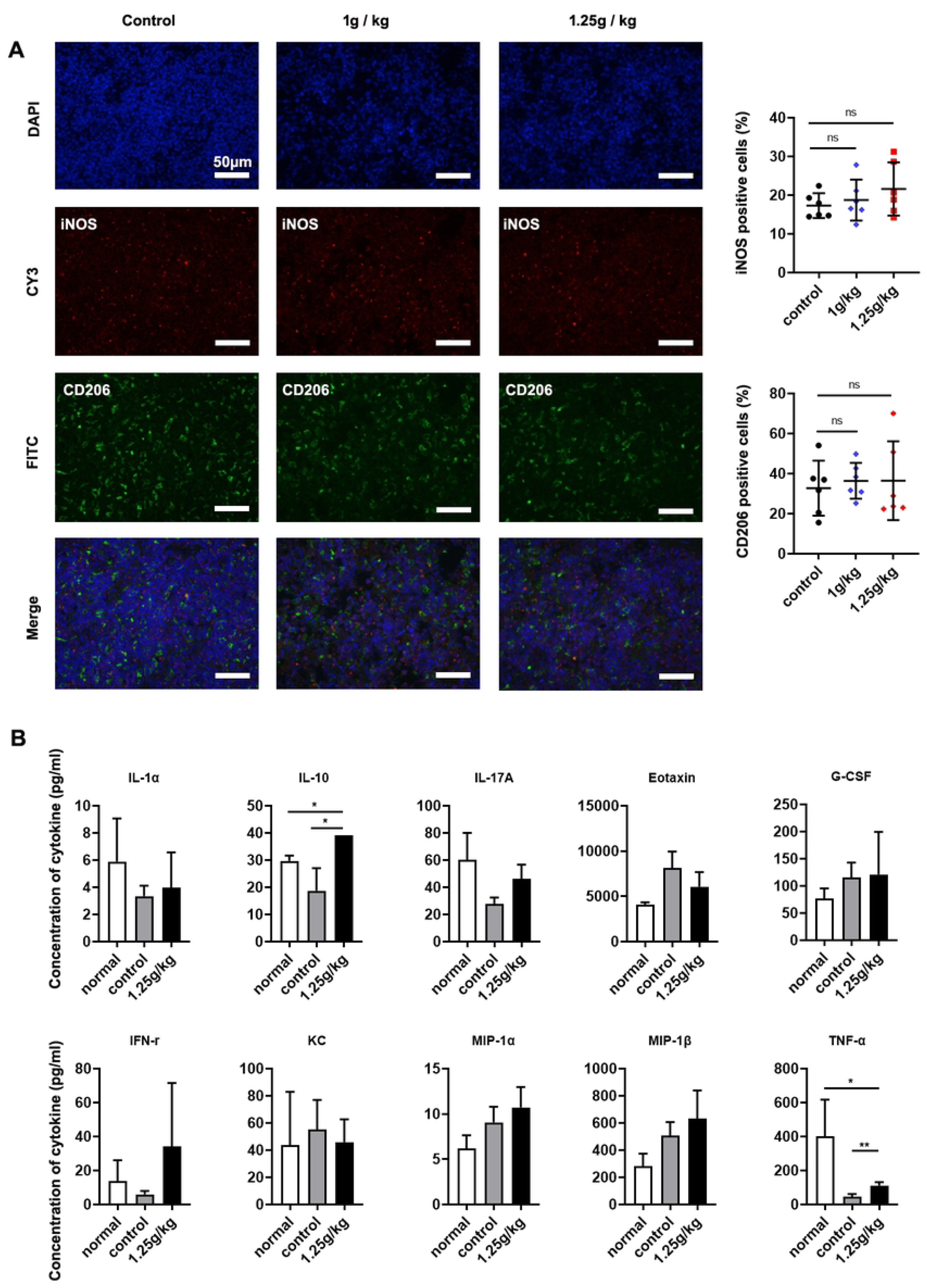
Effect of adding iron supply in *B. microti*-infected mice on macrophage and cytokine secretion. **(A)** Immunofluorescence staining analysis of the expression of M1 and M2 macrophage-specific markers iNOS and CD206 in mouse spleen tissue sections (scale bar = 50 µm; n = 6, per group). Control group, no iron supply in infected mice; 1 or 1.25 g/kg group, subcutaneous injections of 1 or 1.25 g/kg iron dextran to infected mice every two days. **(B)** Expression of IL-1α, IL-10, IL-17A, eotaxin, G-CSF, IFN-γ, KC, MIP-1α MIP-1β, and TNF-α in mouse serum was detected by using Bio-Plex Pro Mouse Cytokine Grp I Panel 23-plex (n = 6, per group). Normal group, uninfected group; control group, infected mice without iron dextran injection; 1.25 g/kg group, infected mice with subcutaneous injections of 1.25 g/kg iron dextran to infected mice every two days. “ns” indicates not significant (p > 0.05), whereas the asterisk indicates significant difference (*, p < 0.05; **, p < 0.01).

### Cytokine analysis

Cytokines in mouse serum were evaluated by Bio-Plex Pro mouse cytokine 23-plex assays. Significant differences were identified for IL-10 and TNF-α (Fig 6B). Other cytokines, including IL-1α, IL-17A, eotaxin, G-CSF, INF-γ, KC, MIP-1α, MIP-1α, MIP-1β, and TNF-α, were not significantly different among the three groups. Based on the results, iron dextran can not change the production of most cytokines in vivo. Although it can stimulate anti-inflammatory IL-10 production, it also stimulates TNF-α production. The result suggested that cytokines in mouse serum were not considerably affected by iron dextran.

## Discussion

*Babesia* parasitizes erythrocytes and digests hemoglobin to utilize the products similar to malaria parasites. However, *Babesia* is not sensitive to ARTs, but malaria parasites are. Moreover, malaria parasites are sensitive to iron chelators [22–27], whereas *Babesia* is not[28]. In particular, malaria parasites can produce and store hemozoin and sustain a high haem level throughout their development in RBCs [9]. Furthermore, *Plasmodium* genome includes all genes relevant to haem synthesis, whereas *Babesia* genomes do not. Apparently, *Babesia* does not rely on haem or iron as much as malaria parasites. Given that ARTs require haem or iron to activate, the requirement of HI likely determines the sensitivity to ARTs. Especially, a recent research proposed a double-kill mechanism of artemisinin against *Plasmodium* through iron-capture and free-radical effects. The theory further suggests that ARTs kill parasites through the interaction of ARTs and HI[12]. Likely, the more a parasite requires, stores, utilizes, or relies on HI, the more it is sensitive to ARTs.

This raises the question of why malaria parasites require far more HI than *Babesia*. Of note, *P. yoelii* 17XNL DNA replication-related genes are expressed more actively than *B. microti*. Moreover, both parasites have the complete PPP enzyme system, but more *P. yoelii* 17XNL showed higher expression than *B. microti*. In particular, *P. yoelii* 17XNL have an HHIP cycle to continuously provided ribose-5-phosphate for nucleotide biosynthesis[12]. If *B. microti* only requires amino acids after the digestion of hemoglobin rather than haem, they likely lack the HHIP cycle. Especially, relevant enzyme genes in PPP and glutathione-related genes in *B.microti* are inactively expressed. Even though they have HHIP cycle, the efficiency of the cycle is not as high as that in *P. yoelii* 17XNL. Clearly, when the HHIP cycle is sustained, DNA synthesis can continuously acquire ribose-5-phosphate, undoubtedly facilitating the production of numerous merozoites. The fact that *P. falciparum* can produce 8-32 merozoites whereas *Babesia* can only produce 2-4 merozoites proves exactly that. Apparently, HI plays a crucial role in the development and reproduction in malaria parasites. Given that malaria parasites sustain a high haem level at nearly all stages in RBCs[9], it suggests that all stages require haem, likely explaining why most stages are sensitive to 24 h of artemether treatment. By contrast, nearly all *B.microti* that expressed DNA synthesis-, antioxidation-, glycolysis-, reproduction- and glutathione-related genes, are not susceptible to artemether. Likely, HI-dependence becomes the Achilles’ heel of malaria parasites under artemisinin action, but it seems to be fortunate to *Babesia*.

To investigate the significance of HI for *Babesia*, we added the iron supply through subcutaneous injections of iron dextran in in vivo experiments. The results showed that the infection rate of *Babesia* increased in a dose-dependent manner. Of note, the increase of the infection rate of *B. microti* is not related to the change of macrophage polarization or cytokine secretion. Apparently, the extra supply of iron is beneficial to promote the reproduction of *B. microti*. Thus, this raises the question of why *Babesia* do not evolve similar mechanisms to malaria parasites to utilize HI to enhance their fertility.

Compared with malaria parasites with an average of more than 20 Mb genomes [29, 30], *Babesia* spp. only have smaller genomes with about 6 Mb [31–33]. In particular, *Babesia* genomes do not include complete haem synthesis enzyme genes. Likely, *Babesia* spp. have not large enough genomes to evolve some similar mechanisms to those in malaria parasites to accumulate and utilize HI, thereby resulting in lower fertility than malaria parasites. However, perhaps fortunately, *Babesia* have no strong HI requirement, no storage of a large amount of HI, and no dependence on HI in all development stages of *Babesia*, making them insusceptible to ARTs attacks.

## Supporting information

**S1 Fig.**
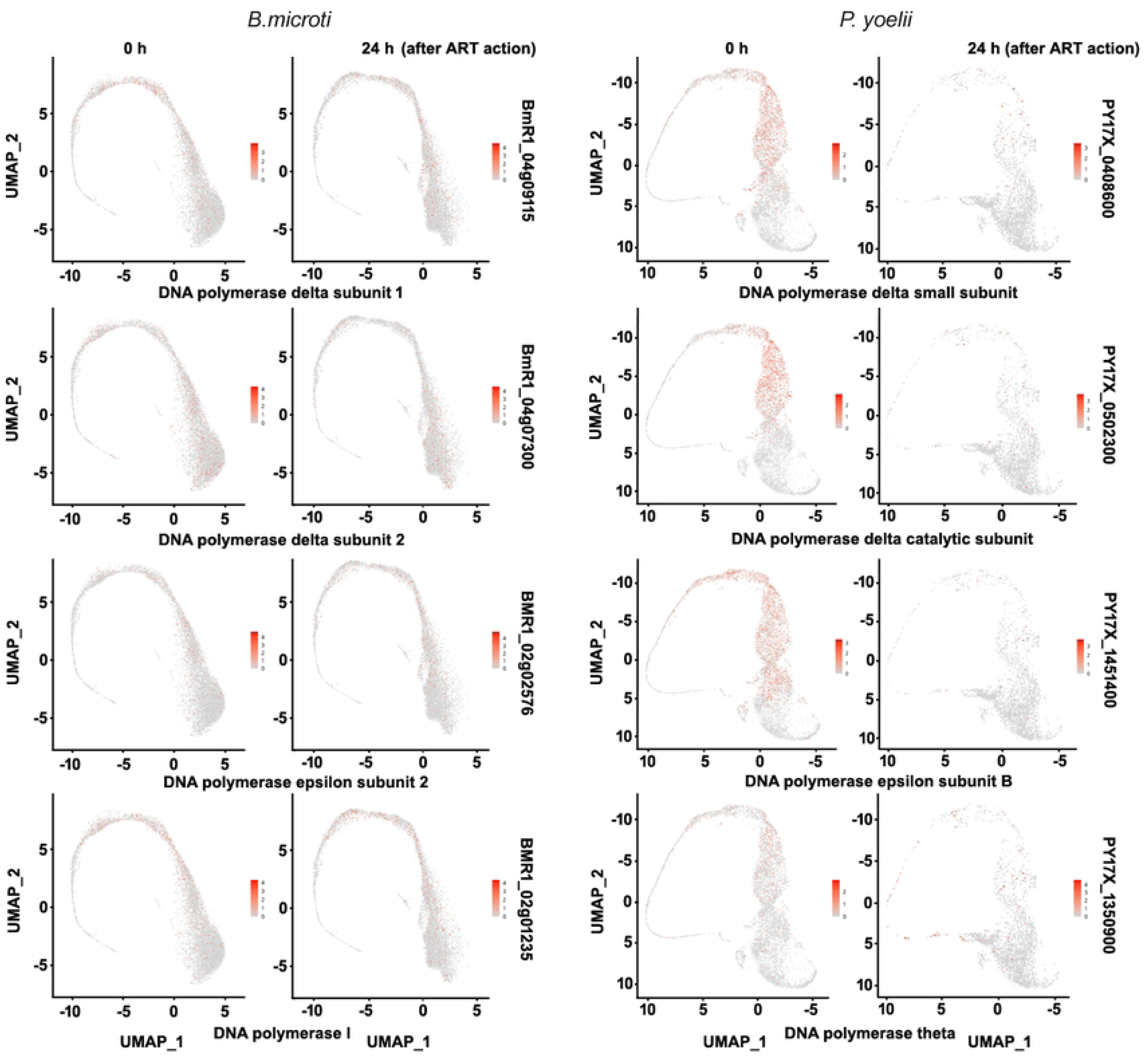
Effect of 24 h of artemether treatment on *B. microti* and *P. yoelii* 17XNL that expressed DNA polymerases. The UMAP plots showed that artemether cannot affect *B. microti* that express DNA polymerase genes, but eliminate *P. yoelii* 17XNL that express similar genes. ART, artemether.

**S1 Table. Comparison of *Babesia* and *Plasmodium* genomes**

**S2 Table. 669 *Plasmodium*-specific genes and GO analysis**

**S3 Table. 924 *Babesia*-specific genes and GO analysis**

**S4 Table. 1591 *Plasmodium-Babesia*-shared genes and GO analysis**

**S5 Table. 1185 *Plasmodium*-specific genes and GO analysis**

**S6 Table. 453 *Babesia*-specific genes and GO analysis**

**S7 Table. 1075 *Plasmodium-Babesia*-shared genes and GO analysis**

**S1 File. Immunofluorescence protocol**

## Conflict of interest

The authors declare that they have no conflict of interest.

## Acknowledgments

I thank LC Sciences (Hangzhou, Zhejiang, China) and Majorbio Co., Ltd (Shanghai, China) for assisting with single-cell RNA sequencing (scRNA-seq) and bioinformatics analysis.

## Funding statement

This research was supported by Innovation Program of Shanghai Municipal Education Commission (201901070007E00017).

## Data availability

All relevant data are within the manuscript.

## Notes

### Competing Interest Statement

The authors have declared no competing interest.

